# *Fate of a fallen ant*: Time-resolved micro-CT analysis of ant digestion in *Nepenthes* pitchers

**DOI:** 10.64898/2025.12.19.695644

**Authors:** Nikitha Sabulal, Gokul Baburaj Sujatha, Anil John Johnson, Sabulal Baby

**Author notes:** **Correspondence** S. Baby, Phytochemistry and Phytopharmacology Division, Jawaharlal Nehru Tropical Botanic Garden and Research Institute, Pacha-Palode, Thiruvananthapuram, Kerala, India. These authors contributed equally to this work (Nikitha Sabulal, Gokul Baburaj Sujatha).

## Abstract

Carnivorous pitcher plants of the genus *Nepenthes* have evolved highly specialized leaf-derived traps that capture and enzymatically digest insect prey to supplement nutrient acquisition in nutrient-deficient habitats. This study aimed to elucidate the structural dynamics underlying prey digestion in *Nepenthes khasiana* using high-resolution micro-computed tomography (micro-CT) integrated with biochemical characterization of the pitcher fluid. Three-dimensional morphometric analysis of the invasive yellow crazy ant, *Anoplolepis gracilipes*, experimentally incubated in *N. khasiana* pitcher fluid, was conducted to assess time-dependent structural degradation. Quantitative micro-CT parameters, including volume, surface area, and surface-to-volume ratio, were analyzed to visualize and quantify time-dependent structural degradation and three-dimensional morphological changes in the prey during digestion. A triphasic digestion sequence was identified - comprising rapid soft-tissue hydrolysis, progressive exoskeletal fragmentation, and stabilization of chitinous remnants. The acidic, enzymatic fluid (pH 2.5-4.0) promoted uniform degradation through synergistic enzymatic, oxidative, and acidification processes, while dissolved CO_2_ enhanced acid stability and catalytic efficiency. The integration of micro-CT morphometrics and biochemical insights reveals a coordinated digestion mechanism in *N. khasiana*, optimized for efficient prey breakdown and nutrient assimilation.

**ONE-SENTENCE SUMMARY:** Micro-CT analysis revealed a triphasic digestion process in *Nepenthes khasiana*, elucidating the structural and enzymatic dynamics of efficient prey degradation.

## INTRODUCTION

*Nepenthes* pitchers are evolutionarily modified leaves that function as passive yet highly efficient traps, employing a combination of physical, chemical, and biological strategies to capture and digest ants and other arthropods. Prey capture in *Nepenthes* involves multiple interacting factors, including peristome geometry and microstructure, toxic nectar, visual and olfactory cues (such as CO_2_, volatiles, fluorescence emissions), and environmental influences (humidity, rainfall) (Bohn & Federle, 2004; Kurup *et al*., 2013; Baby *et al*., 2017; Lathika *et al*., 2025a; Sujatha *et al*., 2026). Once insects lose their footing on the wet peristome, they slide into the viscoelastic digestive fluid at the bottom of the pitcher through the slippery zone (Gaume & Forterre, 2007; Riedel *et al*., 2007), where they undergo enzymatic and microbial degradation (Saganová *et al*., 2018; Bittleston *et al*., 2023). The released nutrients are absorbed through the pitcher walls, while indigestible remains settle at the bottom (Schulze *et al*., 1999).

Among all prey types, ants are the most frequently captured organisms in *Nepenthes* pitchers (Merbach *et al*., 2001; Di Giusto *et al*., 2008; Gaume *et al*., 2016; Lathika *et al*., 2025a; Lathika *et al*., 2025b). Their digestion represents a complex biochemical process involving plant-secreted enzymes and microbial symbionts acting within an acidic pitcher fluid (Rottloff *et al*., 2016; Saganová *et al*., 2018; Gilbert *et al*., 2020). The process begins with the breakdown of the chitinous exoskeleton by chitinases, followed by the degradation of proteins, lipids, and nucleic acids, enabling nutrient absorption through specialized digestive glands on the inner pitcher wall (Rottloff *et al*., 2016).

In our conservatory at the Jawaharlal Nehru Tropical Botanic Garden and Research Institute at Palode in south India, the yellow crazy ant (*Anoplolepis gracilipes* Smith) was observed as a frequent visitor to *Nepenthes khasiana* Hook.f. pitchers (Lee & Yang, 2022; Lathika *et al*., 2025a). Moreover, exoskeletal remnants of ants and other arthropods were often found in senescent pitchers, suggesting variable digestion processes (Fig. 1). Despite in-depth studies on the biochemical aspects of digestion, the intermediate morphological stages of prey degradation within *Nepenthes* pitchers remain poorly understood. Micro-computed tomography (micro-CT) has been extensively applied in insect morphometry to non-destructively visualize and quantify internal and external anatomical structures with high spatial precision (Hita Garcia *et al*., 2017; Jonsson, 2023). This study employs micro-CT to elucidate the sequential morphological transformations of *A. gracilipes* during digestion in *N. khasiana*, aiming to characterize the structural dynamics underlying prey degradation within the pitcher trap.

**Fig. 1.**
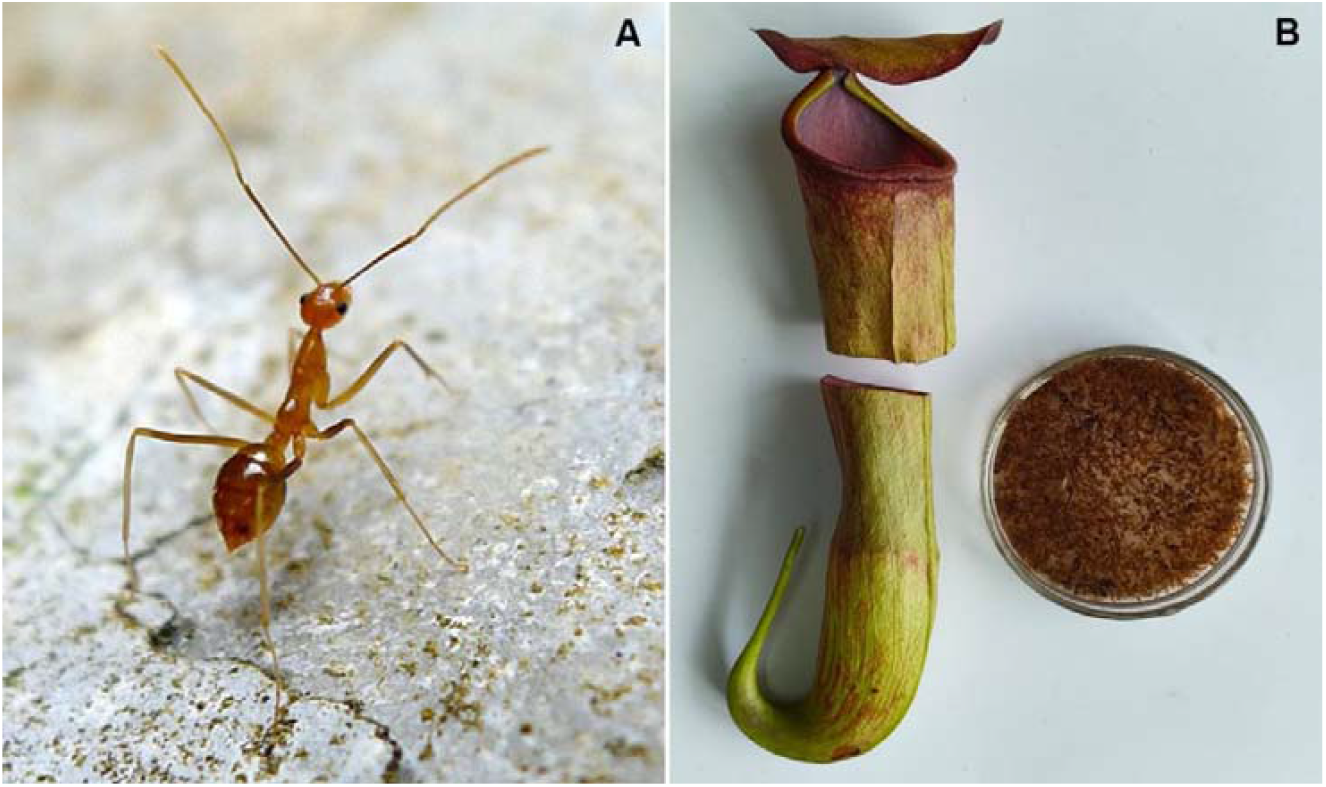
(A) *Anoplolepis gracilipes* ant and (B) digested debris of preys (ants) in *N. khasiana* pitcher fluid.

## MATERIALS AND METHODS

### Ant digestion in *N. khasiana* pitchers

A few selected *N. khasiana* pitchers (Fig. 1) maintained at the conservatory of the Jawaharlal Nehru Tropical Botanic Garden and Research Institute (08°45′00.04″ N, 77°01′41.09″ E) were covered with fine nylon nets prior to opening to prevent prey entry. Upon opening, workers of *A. gracilipes* (yellow crazy ant; Fig. 1), a long-legged ant species commonly visiting *N. khasiana* pitchers, collected from the same native habitat, were gently guided onto the peristomes until they fell into the pitcher fluid. The pitchers were then re-covered with nylon nets to exclude further prey entry. The ants experimentally incubated in pitcher fluids were retrieved from labelled pitchers after 2, 5, and 10 days of digestion (n = 4 per time point). Pitcher fluids containing ants at various digestion stages were transferred to Petri dishes, the fluid volumes were reduced using micropipettes, and the ants were isolated under a handheld lens with an external light source. Control *A. gracilipes* ants (n = 4) were collected from the surrounding vegetation.

All ants were fixed overnight at room temperature in 10% neutral buffered formalin (pH 7.4) containing approximately 0.5% Triton X-100 to facilitate cuticular permeation. After fixation, the samples were washed in neutral buffered formalin and dehydrated sequentially in 30%, 50%, and 70% ethanol (diluted with double-distilled water). Both test (experimental, without pitcher fluid) and control ants were transferred into 70% ethanol, stored in sealed containers, and maintained at 4°C until micro-CT analysis.

### Micro-computed tomography scanning and reconstruction

*Anoplolepis gracilipes* control and test ants (n = 4 each) were imaged using a high-resolution micro-CT scanner (SkyScan 1272, Bruker, Kontich, Belgium) equipped with a Hamamatsu L11871-20 microfocus X-ray source and a XIMEA xiRAY16 detector (7.4 µm pixel size, X/Y ratio = 1.0008). Scans were performed at a source voltage of 40 kV and current of 175 µA, with an exposure time of 583 ms per projection. Frame averaging (2x) was applied to reduce noise, and flat-field correction was enabled with an updating interval of 108 images. No physical filter was used during image acquisition.

The ant specimens were positioned 52.9 mm from the source (camera-to-source distance: 224.2 mm). A total of 750 projection images were acquired over a full 360° rotation using a 0.48° rotation step and step-and-shoot detector motion. The resulting raw images were collected at an effective isotropic voxel size of 3.50 µm and saved in 16-bit TIFF format. Reconstruction was carried out using NRecon software (v 1.7.4.6) with GPU-based processing (GPU Recon Server v 1.7.4). A total of 496 cross-sectional slices were reconstructed from section 286 to 1277, producing PNG images with a matrix size of 872 × 872 pixels and a final voxel size of 7.00 µm. A Hamming filter (cutoff = 100% of Nyquist, α = 0.54) was applied, with ring artifact correction set to 10. Post-alignment correction of 3.5° was performed. Beam hardening correction and HU calibration were not applied. Reconstruction was restricted to a circular region of interest (ROI: top = 1943, bottom = 195, left = 350, right = 2098 pixels). The total reconstruction time was 149 s (0.30 s per slice). The reconstructed dataset was exported as compressed PNG images for subsequent quantitative and qualitative analysis.

### Data analysis

Reconstructed datasets were imported into Avizo 3D (Thermo Fisher Scientific, USA) for three-dimensional visualization, segmentation, and rendering. Morphometric parameters, including head width, thorax length, gaster volume, and appendage dimensions, were quantified using the software’s measurement module. Volume renderings were generated to compare internal anatomical features between control and test ants. Segmentation of exoskeletal and internal structures was performed using a combination of threshold-based selection and manual refinement. For qualitative analysis, 3D models were rotated, sliced, and examined to visualize structural alterations. Quantitative datasets were exported in CSV format for statistical evaluation in R (v 4.3.2).

## RESULTS AND DISCUSSION

High-resolution micro-computed tomography (micro-CT) revealed a progressive, time-dependent degradation of *A. gracilipes* ants exposed to the pitcher fluid of *N. khasiana* over ten days (Fig. 2). Control ants (day 0) exhibited intact morphology with distinct segmentation of the head, thorax and gaster, well-preserved appendages, and homogeneous internal density. The exoskeleton displayed strong X-ray absorption, confirming structural integrity and effective fixation prior to scanning (Fig. S1). After two days of exposure, early signs of digestion appeared as a reduction in internal tissue contrast and thinning of the exoskeleton, particularly at intersegmental joints. Slight softening of the thoracic and abdominal regions indicated enzymatic infiltration of pitcher fluid into internal cavities, although no external fragmentation was yet visible (Figs. 2, S1).

**Fig. 2.**
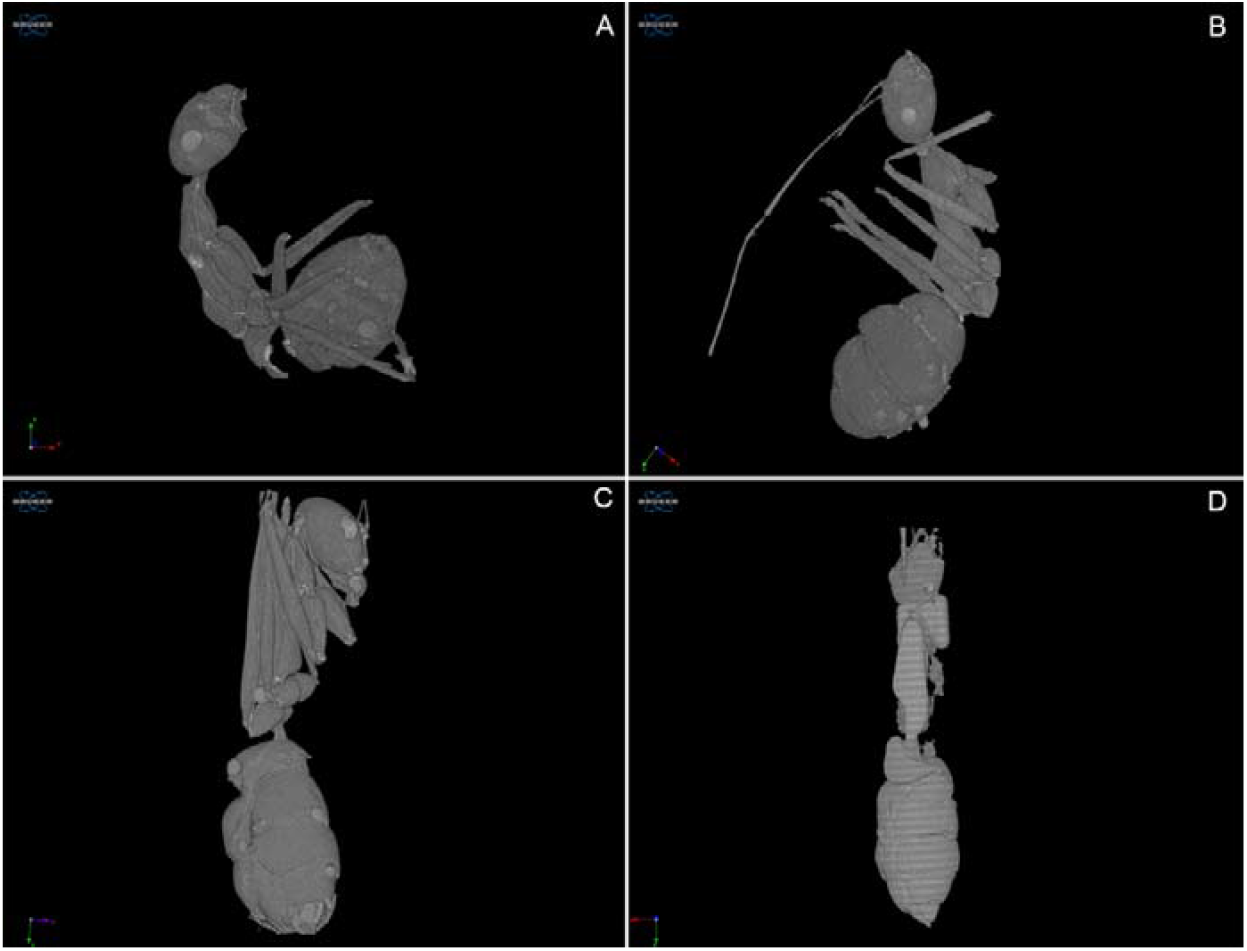
Micro-CT images of progressive breakdown of an *A. gracilipes* ant over 10 days in *N. khasiana* pitcher fluid. (A) Control ant (day 0, no digestion); (B) day 2; (C) day 5; and (D) day 10 of digestion.

By day 5, micro-CT reconstructions showed pronounced structural collapse and fragmentation of the thorax and gaster. Internal voids indicated substantial tissue liquefaction and muscular degradation, and appendages appeared partially disarticulated. Quantitative morphometry revealed an approximate 30-40% reduction in gaster volume and 20-25% reduction in thorax length relative to controls (Fig. S1). After ten days of digestion, only low-density exoskeletal fragments remained (Figs. 1, 2). The thorax and gaster were completely collapsed, with loss of internal tissue and appendages, leaving porous, irregular chitinous residues that represented indigestible remnants. Voxel intensity profiles confirmed near-total loss of soft-tissue density, consistent with advanced digestion and decomposition (Figs. 1, 2, S1, videos S1-S4).

Morphometric analysis demonstrated clear temporal trends in structural degradation (Table 1). The number of layers increased significantly at day 2 (659 ± 91.46) compared with the control (469.25 ± 108.93; *p* < 0.05), but returned near baseline by day 10 (480.75 ± 212.02). The lower vertical position decreased by over 50% at day 2 (0.54 ± 0.27 *vs*. 1.14 ± 0.59; *p* < 0.05), while the upper position slightly increased. Total Volume of Interest (VOI) and object volumes nearly doubled at day 2 (2.19 ± 0.20 *vs*. 1.13 ± 0.28; *p* < 0.01) due to osmotic swelling and fluid penetration, then declined to 1.36 ± 0.63 by day 10. Object surface area followed a similar trend, peaking at day 2 (22.28 ± 1.46; *p* < 0.05) before decreasing by day 10 (14.45 ± 7.11). The surface-to-volume ratio, an indicator of surface complexity, was lowest at day 2 (10.22 ± 0.64 vs. 13.33 ± 1.75; *p* < 0.05), indicating surface smoothening during early digestion, and rose modestly by day 5 (11.40 ± 2.26) and day 10 (10.62 ± 3.85), reflecting renewed surface roughness due to fragmentation of the chitinous exoskeleton. Centroid coordinates (x, y, z) fluctuated within narrow ranges across treatments, indicating uniform degradation rather than localized digestion (Table 1).

**Table 1.** Morphometric parameters of *A. gracilipes* ants during digestion in the pitcher fluid of *N. khasiana* over 2, 5, and 10 days.

| *A. gracilipes* | Control | 2 days | 5 days | 10 days |
| --- | --- | --- | --- | --- |
| Number of layers | 469.25 ± 108.93 | 659 ± 91.46 | 534 ± 168.13 | 480.75 ± 212.02 |
| Lower vertical position | 1.14 ± 0.59 | 0.54 ± 0.27 | 1.06 ± 0.77 | 1.31 ± 0.88 |
| Upper vertical position | 4.42 ± 0.38 | 5.15 ± 0.38 | 4.79 ± 0.47 | 4.66 ± 0.67 |
| Pixel size | 7.0 ± 0.00 | 7.0 ± 0.00 | 7.0 ± 0.00 | 7.0 ± 0.00 |
| Lower grey threshold | 90 ± 0.00 | 90 ± 0.00 | 90 ± 0.00 | 90 ± 0.00 |
| Upper grey threshold | 255 ± 0.00 | 255 ± 0.00 | 255 ± 0.00 | 255 ± 0.00 |
| Total VOI volume | 1.13 ± 0.28 | 2.19 ± 0.20 | 1.60 ± 0.59 | 1.36 ± 0.63 |
| Object volume | 1.13 ± 0.28 | 2.19 ± 0.20 | 1.60 ± 0.59 | 1.36 ± 0.63 |
| Percent object volume | 100 ± 0.00 | 100 ± 0.00 | 100 ± 0.00 | 100 ± 0.00 |
| Total VOI surface | 14.89 ± 3.23 | 22.28 ± 1.46 | 17.37 ± 3.31 | 14.45 ± 7.11 |
| Object surface | 14.89 ± 3.23 | 22.28 ± 1.46 | 17.37 ± 3.31 | 14.45 ± 7.11 |
| Intersection surface | 14.89 ± 3.23 | 22.28 ± 1.46 | 17.37 ± 3.31 | 14.45 ± 7.11 |
| Object surface/<br>volume ratio | 13.33 ± 1.75 | 10.22 ± 0.64 | 11.40 ± 2.26 | 10.62 ± 3.85 |
| Object surface density | 13.33 ± 1.75 | 10.22 ± 0.64 | 11.40 ± 2.26 | 10.62 ± 3.85 |
| Centroid (x) | 1.73 ± 0.69 | 1.80 ± 0.79 | 1.55 ± 0.40 | 1.39 ± 0.24 |
| Centroid (y) | 1.81 ± 0.73 | 1.61 ± 0.42 | 2.01 ± 0.89 | 1.33 ± 0.23 |
| Centroid (z) | 2.94 ± 0.32 | 3.15 ± 0.56 | 2.90 ± 0.41 | 3.08 ± 0.58 |

VOI: Volume of Interest; Each data is an average of for measurements (mean ± SD, n = 4).

Sequential micro-CT imaging thus captured a continuous three-phase digestive trajectory within the viscous, acidic fluid of the pitchers of *N. khasiana*, a montane species endemic to Northeast India characterized by waxy inner walls, viscoelastic fluid, and dense distributions of digestive glands that secrete acid-stable hydrolases enabling efficient prey retention and enzymatic breakdown. In the present 10-day ant digestion study, representing approximately one-fifth to one-third of the functional lifespan of a *N. khasiana* pitcher (typically 4-8 weeks under natural conditions) (Baby *et al*., 2017; Devi *et al*., 2019; Dkhar *et al*., 2020), the digestion of *A. gracilipes* ants proceeded through a well-defined temporal sequence: (i) enzymatic infiltration and osmotic swelling (day 0-2), (ii) structural disintegration through proteolytic and chitinolytic degradation (day 3-5), and (iii) residual stabilization of indigestible fragments (day 6-10). The initial infiltration phase was characterized by increased object volume and reduced internal contrast, consistent with early enzymatic activity by acid-stable aspartic proteases such as nepenthesins secreted from the glandular surface (An *et al*., 2000; Hatano & Hamada, 2008; Buch *et al*., 2015). The subsequent disintegration phase involved marked tissue loss and surface simplification, reflecting rapid hydrolysis of soft tissues by proteases and chitinases that jointly facilitate nutrient release (Rottloff *et al*., 2016; Saganová *et al*., 2018). The final phase resulted in nearly complete digestion of organic matter, leaving only chitinous exoskeletal residues that accumulated at the pitcher base, consistent with prior reports that *Nepenthes* pitchers retain partially digested insect cuticles as persistent remnants following nutrient absorption (Fig. 1) (Schulze *et al*., 1999; Renner & Specht, 2013). This sequential 10-day trajectory thus represents a single, complete digestion cycle occurring within the early active period of the pitcher’s lifespan, mirroring the natural rhythm of prey capture, enzymatic digestion, and residue stabilization typical of functional *N. khasiana* pitchers.

These digestion dynamics reflect coordinated enzymatic and physicochemical processes characteristic of *Nepenthes* pitchers, regulated hormonally through jasmonic acid (Buch *et al*., 2015; Yilamujiang *et al*., 2016; Saganová *et al*., 2018). Jasmonate signalling acts as a molecular switch activating digestive pathways following prey capture, paralleling plant defense responses (Mithöfer, 2011; Pavlovič & Mithöfer, 2019). The enzyme-mediated breakdown of proteins, lipids and chitin occurs in an acidic, enzyme-rich fluid (pH 2.5-4.0), which provides optimal conditions for proteolytic and chitinolytic activity while maintaining redox stability (An *et al*., 2000; Rottloff *et al*., 2016).

The physicochemical environment of *N. khasiana* further enhances digestion. Upon prey induction, the plant secretes naphthoquinones such as droserone and 5-*O*-methyl droserone (Raj *et al*., 2011; Lathika *et al*., 2025b), imparting the orange-red hue to the pitcher fluid. These oxidative metabolites, together with plumbagin and (+)-isoshinanolone localized in the peristome and nectar, act as antimicrobial and insecticidal agents that prevent microbial spoilage and aid in prey immobilization (Lathika *et al*., 2025a; Lathika *et al*., 2025b). Despite these inhibitory effects, the pitcher fluid supports a specialized community of acid-tolerant inquilines, including bacteria, protozoa and dipteran larvae, that contribute to secondary digestion through proteolytic, chitinolytic and ammonifying activities (Adlassnig *et al*., 2011; Gilbert *et al*., 2020). These microbial symbionts complement plant-derived enzymes, forming an integrated digestive ecosystem that couples chemical defense, antimicrobial regulation and cooperative enzymatic hydrolysis.

The internal gas composition of *N. khasiana* pitchers also contributes to digestive efficiency. Elevated CO_2_ concentrations (*N. khasiana* unopen pitcher: 4053.76 ± 1188.84 ppm, n = 9; open pitcher 476.75 ± 59.83 ppm, n = 6) reported by Baby *et al*. (2017) not only attract arthropods but also enhance pitcher fluid chemistry. Dissolved CO_2_ equilibrates to carbonic acid, reinforcing acidity and buffering against local pH increases during prey decomposition (An *et al*., 2000). Continuous CO_2_ release from pitcher tissues maintains autogenic acidification even before prey capture, ensuring digestive readiness.

Ants represent the most frequent and nutritionally valuable prey in *Nepenthes* pitchers (Merbach *et al*., 2001; Di Giusto *et al*., 2008; Gaume *et al*., 2016; Lathika *et al*., 2025a; 2025b). Their high protein content (35-60% dry weight; Rumpold & Schlüter, 2013; Hasnan *et al*., 2023) provides a major nitrogen source in nutrient-poor montane environments. The invasive ant *A. gracilipes*, characterized by rapid movement and long-legged morphology, was the predominant prey species observed in *N. khasiana* at Palode, Kerala. Similar dominance of this ant species has been reported in *N. weda* (Mansur & Brearley, 2024) and *N. rafflesiana* (Gaume & Di Giusto, 2009), underscoring its ecological importance across diverse *Nepenthes* habitats.

The observed digestion pattern aligns with previous studies showing that prey capture and degradation in *Nepenthes* involve multiple interacting factors, including peristome microstructure, toxic nectar, visual and olfactory cues such as CO_2_ and fluorescence (Bohn & Federle, 2004; Kurup *et al*., 2013; Baby *et al*., 2017; Lathika *et al*., 2025a), and environmental influences such as humidity and rainfall (Gaume & Forterre, 2007; Riedel *et al*., 2007). These mechanisms, combined with enzymatic hydrolysis, naphthoquinone-mediated defense, CO_2_-assisted buffering and microbial symbiosis, exemplify a finely tuned digestive system in *N. khasiana*. Overall, the integration of acidic enzymatic hydrolysis, oxidative metabolites, hormonal regulation, CO_2_ buffering and microbial cooperation enables *N. khasiana* to efficiently degrade both soft and sclerotized prey tissues. This synergistic system maximizes nutrient recovery from ant prey, explaining the ecological success of *N. khasiana* in oligotrophic habitats. Sequential micro-CT analysis provided non-destructive, quantitative evidence linking morphological transformation with enzyme-driven digestion, thereby elucidating the mechanistic basis of prey decomposition and nutrient recycling in carnivorous plants.

## CONCLUSIONS

This study demonstrates that *N. khasiana* digests the invasive yellow crazy ant (*A. gracilipes*) through a distinct, three-phase sequence involving enzymatic hydrolysis of soft tissues, progressive exoskeletal fragmentation, and stabilization of indigestible chitinous residues. Sequential micro-CT imaging provided non-destructive, quantitative evidence of these transformations, confirming uniform digestion across the prey body and revealing coordinated proteolytic, chitinolytic, and osmotic activities within the acidic pitcher fluid. In combination with these enzymatic mechanisms, neurotoxic and antimicrobial naphthoquinones and a CO_2_-enriched acidic milieu enhance digestive efficiency and maintain microbial stability. Together, these biochemical and physicochemical processes underline the adaptive optimization of *N. khasiana*’s digestive physiology for nutrient extraction from hard-bodied prey. The frequent capture of *A. gracilipes* further highlights its ecological significance as a major nitrogen source, linking invasive ant dynamics with nutrient acquisition in *Nepenthes* habitats.

## AUTHOR CONTRIBUTIONS

Conceptualization: N.S., G.B.S., S.B.; methodology, formal analysis and data acquisition: N.S., G.B.S., A.J.J., S.B.; fund acquisition: S.B.; writing - original manuscript: N.S., G.B.S., A.J.J., S.B.; writing - review, editing: N.S., G.B.S., A.J.J., S.B.

## ACKNOWLEDGEMENTS

We acknowledge Garden Management Division, Jawaharlal Nehru Tropical Botanic Garden and Research Institute, Palode for facilitating field studies and providing us the plant specimens.

## SUPPORTING INFORMATION

Additional supporting information may be found online in the Supporting Information section at the end of the article.

**Figure S1**. Time-resolved micro-CT visualization of the yellow crazy ant, *Anoplolepis gracilipes*, digestion within the pitcher fluid of *Nepenthes khasiana*. Three-dimensional reconstructions illustrate progressive structural degradation of *A. gracilipes*, experimentally incubated in the acidic, enzyme-rich pitcher fluid over 10 days. Groups 1-4 correspond to independent projection datasets acquired from different viewing angles (Group 1: Image 000, Group 2: 300, Group 3: 600, & Group 4: 749), each comprising four time points: (A) control, (B) day 2, (C) day 5, and (D) day 10.

**Videos S1-S4**: Three-dimensional video reconstructions showing the progressive degradation of the ant, *A. gracilipes*, incubated in the pitcher fluid of *N. khasiana* over a 10-day period. Video S1: control ant; video S2: day 2 digestion; video S3: day 5; video S4: day 10, illustrating the sequential disintegration within the digestive fluid. 3D videos were reconstructed from micro-CT image stacks using 3D Slicer ver 5.8.1.

