## Supplementary material for "*Fate of a fallen ant*: Time-resolved micro-CT analysis of ant digestion in *Nepenthes* pitchers": Figure S1

**Correspondence**

S. Baby

Phytochemistry and Phytopharmacology Division

Jawaharlal Nehru Tropical Botanic Garden and Research Institute

Pacha-Palode, Thiruvananthapuram, Kerala, India.

^†^ These authors contributed equally to this work.

**SHORT TITLE:** Ant digestion in *Nepenthes* pitchers.

**ONE-SENTENCE SUMMARY:** Micro-CT analysis revealed a triphasic digestion process in *Nepenthes khasiana*, elucidating the structural and enzymatic dynamics of efficient prey degradation.

**ABSTRACT**

Carnivorous pitcher plants of the genus *Nepenthes* have evolved highly specialized leaf-derived traps that capture and enzymatically digest insect prey to supplement nutrient acquisition in nutrient-deficient habitats. This study aimed to elucidate the structural dynamics underlying prey digestion in *Nepenthes khasiana* using high-resolution micro-computed tomography (micro-CT) integrated with biochemical characterization of the pitcher fluid. Three-dimensional morphometric analysis of the invasive yellow crazy ant, *Anoplolepis gracilipes*, experimentally incubated in *N. khasiana* pitcher fluid, was conducted to assess time-dependent structural degradation. Quantitative micro-CT parameters, including volume, surface area, and surface-to-volume ratio, were analyzed to visualize and quantify time-dependent structural degradation and three-dimensional morphological changes in the prey during digestion. A triphasic digestion sequence was identified - comprising rapid soft-tissue hydrolysis, progressive exoskeletal fragmentation, and stabilization of chitinous remnants. The acidic, enzymatic fluid (pH 2.5-4.0) promoted uniform degradation through synergistic enzymatic, oxidative, and acidification processes, while dissolved CO₂ enhanced acid stability and catalytic efficiency. The integration of micro-CT morphometrics and biochemical insights reveals a coordinated digestion mechanism in *N. khasiana*, optimized for efficient prey breakdown and nutrient assimilation.

*Keywords*: *Nepenthes khasiana*; *Anoplolepis gracilipes*; prey digestion; micro-CT; morphometry; enzymatic degradation.

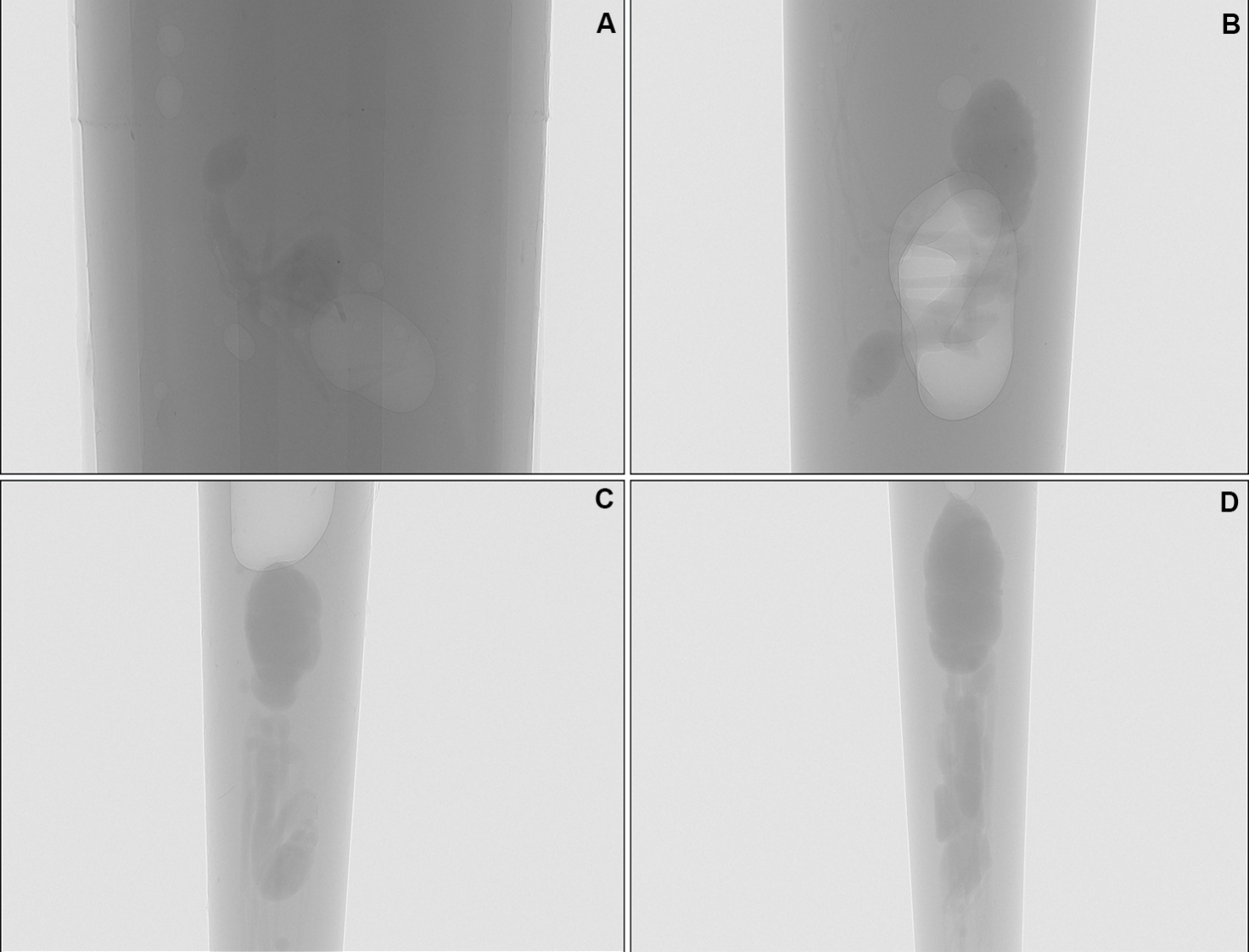

Group 1

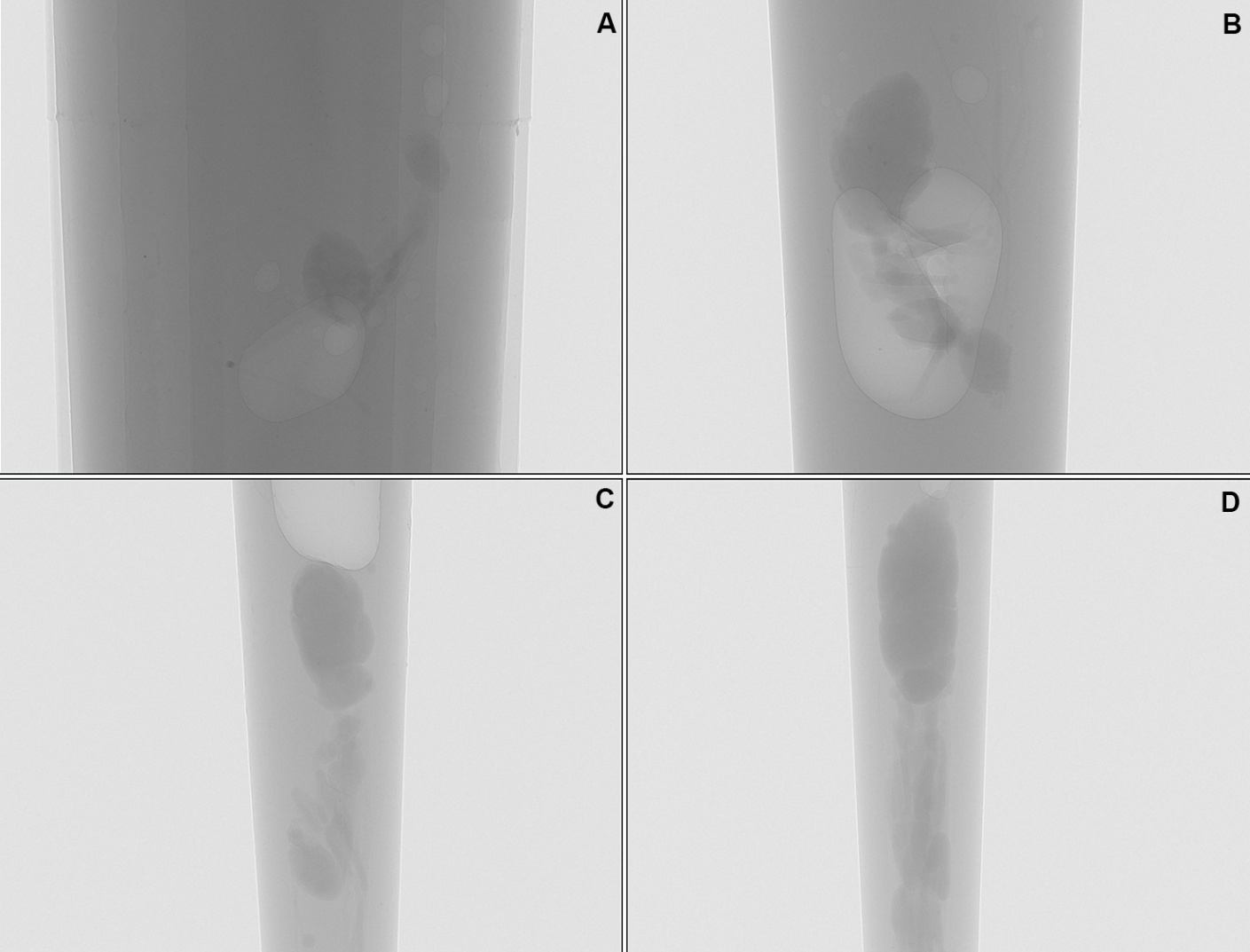

Group 2

Group 3

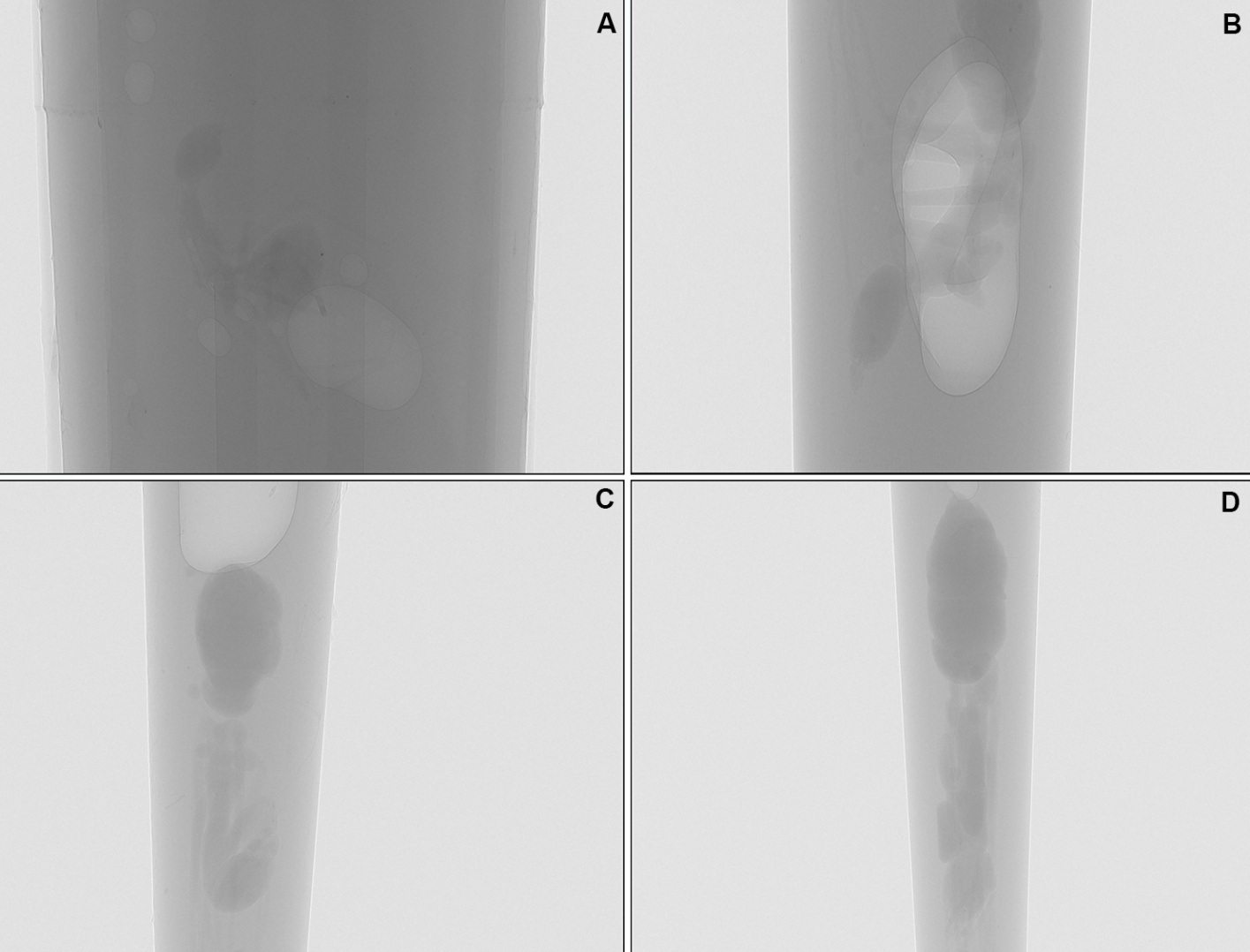

Group 4

**Figure S1.** Time-resolved micro-CT visualization of the yellow crazy ant, *Anoplolepis gracilipes*, digestion within the pitcher fluid of *Nepenthes khasiana*.
Three-dimensional reconstructions illustrate progressive structural degradation of *A. gracilipes*, experimentally incubated in the acidic, enzyme-rich pitcher fluid over 10 days. Groups 1-4 correspond to independent projection datasets acquired from different viewing angles (Group 1: Image 000, Group 2: 300, Group 3: 600, & Group 4: 749), each comprising four time points: (A) control, (B) day 2, (C) day 5, and (D) day 10. Early soft-tissue hydrolysis and cuticular thinning were evident by Day 2, followed by pronounced exoskeletal fragmentation and internal collapse by Day 5. By Day 10, only partially eroded chitinous remnants remained, indicating stabilization of digestion-resistant fragments. Specimens were scanned using a Bruker SkyScan 1272 micro-CT system (40 kV, 175 µA; voxel size 3.5 µm); scale bars 500 µm.

**Videos S1-S4**: Three-dimensional video reconstructions showing the progressive degradation of the ant, *A. gracilipes*, incubated in the pitcher fluid of *N. khasiana* over a 10-day period. Video S1: control ant; video S2: day 2 digestion; video S3: day 5; video S4: day 10, illustrating the sequential disintegration within the digestive fluid. 3D videos were reconstructed from micro-CT image stacks using 3D Slicer ver 5.8.1.
